# Phosphorylation of PSD-95 at Serine 73 in dCA1 is required for extinction of contextual fear

**DOI:** 10.1101/2023.01.11.523603

**Authors:** Magdalena Ziółkowska, Malgorzata Borczyk, Anna Cały, Maria Nalberczak-Skóra, Agata Nowacka, Małgorzata Alicja Śliwińska, Kacper Łukasiewicz, Edyta Skonieczna, Kamil F. Tomaszewski, Tomasz Wójtowicz, Jakub Włodarczyk, Tytus Bernaś, Ahmad Salamian, Kasia Radwanska

**Affiliations:** Laboratory of Molecular Basis of Behavior, the Nencki Institute of Experimental Biology of Polish Academy of Sciences; Department Molecular Neuropharmacology, Maj Institute of Pharmacology of Polish Academy of Sciences, Warsaw, Poland; Laboratory of Imaging Tissue Structure and Function, The Nencki Institute of Experimental Biology of Polish Academy of Sciences, Warsaw, Poland; Laboratory of Cell Biophysics, the Nencki Institute of Experimental Biology of Polish Academy of Sciences; Department of Anatomy and Neurology, VCU School of Medicine, 1101 East Marshall Street, Richmond, Virginia 23298

**Keywords:** PSD-95, fear extinction, dorsal CA1, synaptic plasticity, hippocampus, αCaMKII

## Abstract

The updating of contextual memories is essential for survival in a changing environment. Accumulating data indicate that the dorsal CA1 area (dCA1) contributes to this process. However, the cellular and molecular mechanisms of contextual fear memory updating remain poorly understood. Postsynaptic density protein 95 (PSD-95) regulates the structure and function of glutamatergic synapses. Here, using dCA1-targeted genetic manipulations *in vivo*, combined with *ex vivo* 3D electron microscopy and electrophysiology, we identify a novel, synaptic mechanism that is induced during attenuation of contextual fear memories and involves phosphorylation of PSD-95 at Serine 73 in dCA1. Our data provide the proof that PSD-95-dependent synaptic plasticity in dCA1 is required for updating of contextual fear memory.

## INTRODUCTION

The ability to form, store, and update memories is essential for animal survival. In mammals, the formation, recall and updating of memories involve the hippocampus (Frankland and Bontempi, 2005; Neves et al., 2008; Baldi and Bucherelli, 2015). In particular, formation of memories strengthens the Schaffer collateral-to-dorsal CA1 area (dCA1) synapses through N-methyl-D-aspartate receptor (NMDAR)-dependent forms of synaptic plasticity (Bliss and Collingridge, 1993; Morris et al., 2003; Abraham et al., 2019) linked with growth and addition of new dendritic spines (harbouring glutamatergic synapses) (Restivo et al., 2009; Radwanska et al., 2011; Mahmmoud et al., 2015; Aziz et al., 2019). Although some studies also found long-term depression of synaptic transmission during hippocampal-dependent tasks (Kemp and Manahan-Vaughan, 2007; Goh and Manahan-Vaughan, 2013). Similarly, updating and extinction of memories induces functional, structural, and molecular alterations of dCA1 synapses (Garín-Aguilar et al., 2012; Stansley et al., 2018; Schuette et al., 2020). Accordingly, NMDAR-dependent plasticity of dCA1 synapses is commonly believed to be a primary cellular learning mechanism. Surprisingly, the role of dCA1 synaptic plasticity in memory formation has been recently questioned. Local genetic manipulations that impair synaptic function and plasticity specifically in dCA1 affect spatial choice and incorporation of salience information into cognitive representations, rather than formation of cognitive maps and memory engrams (Bannerman et al., 2012, 2014; Hirsch et al., 2015; Cały et al., 2021; Kaganovsky et al., 2022). On the other hand, the role of dCA1 synaptic plasticity in the updating and extinction of existing hippocampus-dependent memories has not been tested yet. Understanding the molecular and cellular mechanisms that underlie fear extinction memory is crucial to develop new therapeutic approaches to alleviate persistent and unmalleable fear memories.

PSD-95 is the major scaffolding protein of glutamatergic synapses (Cheng et al., 2006). It directly interacts with NMDARs and with AMPARs through an auxiliary protein, stargazin (Kornau et al., 1995; Schnell et al., 2002). Interaction of PSD-95 with stargazin regulates the synaptic content of AMPARs (Chetkovich et al., 2002; Schnell et al., 2002; Bats et al., 2007). Accordingly, PSD-95 affects stability and maturation as well as functional and structural plasticity of glutamatergic synapses (Migaud et al., 1998; Béïque and Andrade, 2003; Stein et al., 2003; Ehrlich and Malinow, 2004; Ehrlich et al., 2007; Nikonenko et al., 2008; Steiner et al., 2008; Sturgill et al., 2009; Chen et al., 2011; Taft and Turrigiano, 2014). Synaptic localisation of PSD-95 is controlled by a range of posttranslational modifications with opposing effects on its synaptic retention as well as synaptic function and plasticity (Vallejo et al., 2017). Here, in order to test the role of dCA1 excitatory synapses in extinction of fear memories, we focused on phosphorylation of PSD-95 at Serine 73 (S73). Phosphorylation of PSD-95(S73) enables PSD-95 dissociation from the complex with GluN2B, and its trafficking to terminate synaptic growth after stimulation of NMDA receptors, and is necessary for PSD-95 protein downregulation during NMDAR-dependent long-term depression of synaptic transmission (LTD) (Steiner et al., 2008; Nowacka et al., 2020). PSD-95(S73) is phosphorylated by the calcium and calmodulin-dependent kinase II (CaMKII) (Gardoni et al., 2006; Steiner et al., 2008). Importantly, both authophosphorylation-deficient αCaMKII mutant mice (αCaMKII-T286A) (Giese et al., 1998) and the loss-of-function PSD-95 mutants lacking the guanylate kinase domain of PSD-95 (Migaud et al., 1998) show impaired extinction of contextual fear (Radwanska et al., 2011; Fitzgerald et al., 2015), suggesting that αCaMKII and PSD-95 interact to regulate contextual fear extinction.

The present study tests the role of PSD-95(S73) phosphorylation in the dorsal hippocampus in fear memory extinction by integrated analyses of PSD-95 protein expression and phosphorylation, dCA1-targeted expression of phosphorylation-deficient PSD-95 protein (with serine 73 mutated to alanine, S73A) as well as examination of dendritic spines morphology with nanoscale resolution enabled by electron microscopy. We show that phosphorylation of PSD-95(S73) is necessary for contextual fear extinction-induced PSD-95 protein regulation and remodelling of glutamatergic synapses. Moreover, it is not necessary for fear memory formation but required for fear extinction even after extensive fear extinction training. Overall, our data show for the first time that the dCA1 PSD-95(S73) phosphorylation is required for extinction of the contextual fear memory.

## RESULTS

### The contextual fear extinction affects PSD-95 protein levels and morphology of dendritic spines in dCA1

To investigate the role of dCA1 excitatory synapses in contextual fear memory extinction, we trained Thy1-GFP(M) mice (that allow for visualisation of dendritic spines) (Feng et al., 2000) in contextual fear conditioning (CFC). The animals showed low freezing levels in the novel context before delivery of 5 electric shocks (US), after which the freezing levels increased during the rest of the training session (**Figure** 1A). Twenty-four hours later, one group of mice was sacrificed (5US) (mice were randomly assigned to the experimental groups), and the second group was re-exposed to the training context for 30 minutes without presentation of US for extinction of contextual fear (Ext). Freezing levels were high at the beginning of the session and decreased within the session indicating formation of fear extinction memory (t = 3.720, df = 6, P < 0.001). Mice were sacrificed immediately after the fear extinction session. Twenty-four hours later, the third group of mice was re-exposed to the training context (without US) to test consolidation of fear extinction memory (test). Freezing levels were lower during the test as compared to the beginning of the extinction, indicating that our protocol resulted in efficient formation of long-term contextual fear extinction memory (P = 0.026). The mouse brains were sliced, the brain sections immunostained to detect PSD-95 protein using specific antibodies and imaged with a confocal microscope. The scans were analysed to asses PSD-95 protein levels [linear density of PSD-95-positive puncta (PSD-95^+^) per 1 μm of a dendrite and mean grey value of PSD-95^+^ per dendritic spine] and dendritic spines linear density and area [enabled by Thy1-GFP(M) transgene] (**Figure** 1B-C). As dendritic spines change in dCA1 after CFC in a dendrite-specific manner (Restivo et al., 2009), the expression of PSD-95 protein, and its colocalization with dendritic protrusions, were analysed in three domains of dCA1: stratum oriens (stOri), stratum radiatum (stRad) and stratum lacunosum-moleculare (stLM) (**Figure** 1D-F).

**Figure 1.**
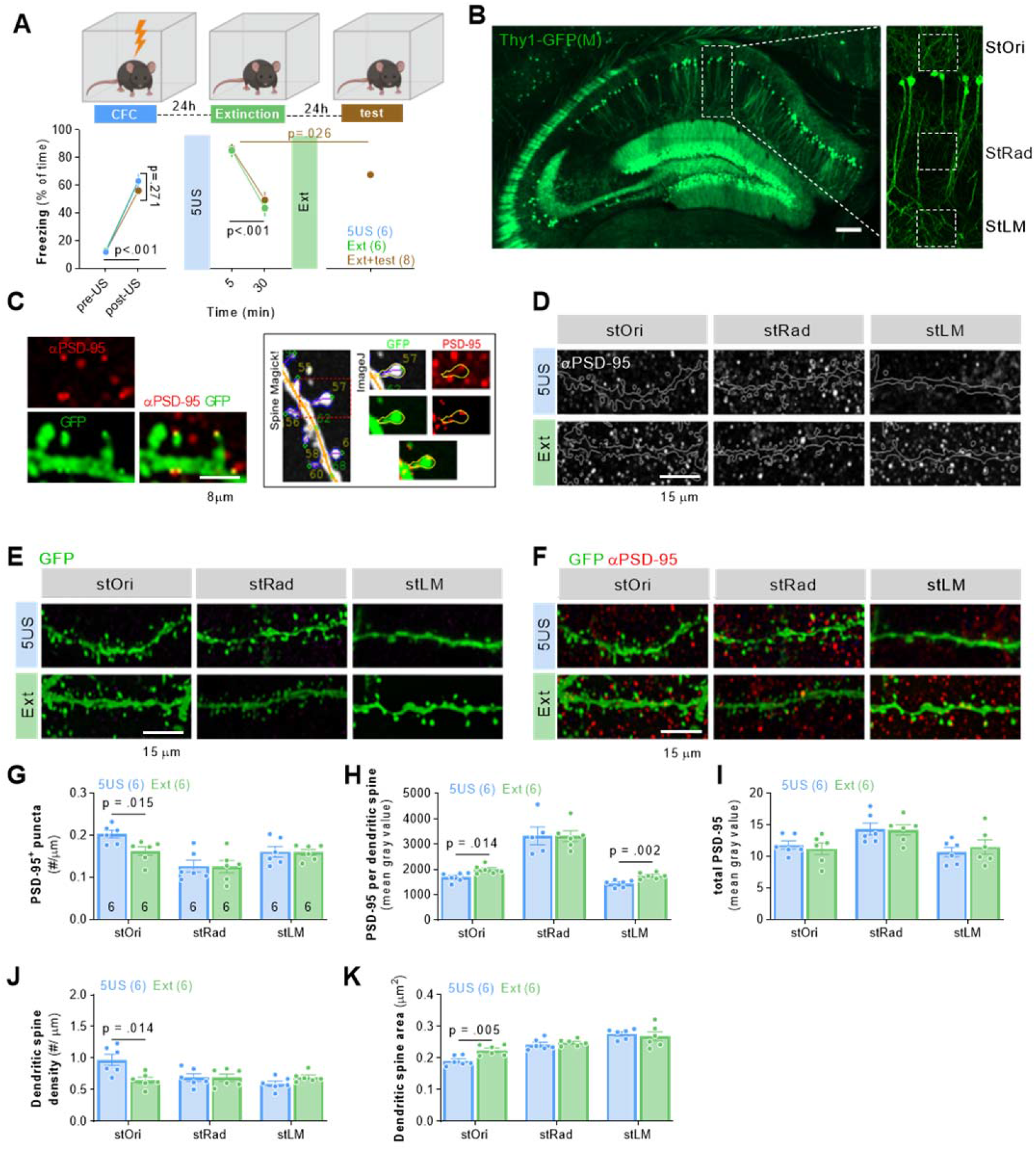
Extinction of contextual fear memory regulates PSD-95 protein levels and remodelling of dendritic spines in dCA1. **(A)** Experimental timeline and freezing levels during training. Mice underwent CFC and were sacrificed 24 hours later (5US, n = 6) or after re-exposure to the training context without electric shocks (Ext, n = 6) (two-way repeated-measures ANOVA, effect of training: F(1, 10) = 77.86, P < 0.0001). **(B-C)** Dendritic spines and PSD-95 expression were analysed in three domains of the dendritic tree of dCA1 pyramidal neurons (stOri, stRad and stLM) in Thy1-GFP(M) male mice. **(B)** Microphotography of dCA1 and dendritic tree domains. **(C)** High magnification of confocal scans showing colocalization of PSD-95 immunostaining and dendritic spines, and the analysis in SpineMagick! and ImageJ. **(D-F)** Representative confocal images (maximum projections of z-stacks composed of 20 scans) of PSD-95 immunostaining, GFP and their colocalization are shown for three domains of dCA1. Scale bar, 15 μm. **(G-I)** Summary of data showing density of PSD-95^+^ puncta (two-way repeated-measures ANOVA with Tukey’s multiple comparisons test (marked on the graphs), effect of training: F(2, 13) = 1.30, P = 0.305), PSD-95 expression per dendritic spine (effect of training: F(2, 15) = 5.653, P = 0.015) and total PSD-95 expression (effect of training: F(2, 14) = 1.126, P = 0.3521). **(J-K)** Summary of data showing dendritic spine density (effect of training: F(2, 44) = 2.851, P = 0.069; a region effect: F(1.983, 43.63) = 6.293, P = 0.004; training × region interaction: F(4, 44) = 5.389, P = 0.001) and average dendritic spine area (two-way repeated-measures ANOVA with Tukey’s multiple comparisons test; effect of training: F(2, 42) = 1.630, P = 0.208; a region effect: F(2, 42) = 46.49, P < 0.001; training × region interaction: F(4, 42) = 2.121, P = 0.095). The analyses were conducted in stOri (mouse/dendrite/spine: 5US = 6/25/650; Ext = 6/37/925). For G-I, each dot represents one mouse. For G-J, M means ± SEM are shown. For K, medians ± IQR are shown.

The analysis of the confocal scans revealed that there were less PSD-95^+^ puncta after fear extinction, as compared to the 5US group in stOri, but not in other dCA1 strata (**Figure** 1G). There was also a significant effect of the training on PSD-95 protein levels per dendritic spine. In the stOri and stLM, PSD-95 levels per dendritic spine increased after extinction, as compared to the 5US group (**Figure** 1H). No difference in PSD-95 levels per dendritic spine was observed between the groups in stRad. Interestingly, when total PSD-95 levels were analysed (as mean grey value of microphotographs) we found no differences between the experimental groups in three strata of dCA1 (**Figure** 1I), indicating bidirectional PSD-95 changes (elimination of PSD-95^+^ puncta and increased intensity of the remaining puncta).

Next, we checked whether the changes in PSD-95 protein levels were associated with dendritic spine remodelling. In stOri, dendritic spine density decreased after extinction as compared to the 5US mice (**Figure** 1J). No changes in dendritic spine density were observed in the stRad and stLM. Moreover, the median dendritic spine area was increased in stOri after extinction, compared to the 5US group, resembling the changes of PSD-95 protein levels. No changes in the median dendritic spine area were observed in the stRad and stLM (**Figure** 1K). In a separate experiment we found that these dendritic spine changes were transient, as they were not observed 60 minutes after contextual fear extinction session, and they were specific for fear extinction, as we did not find such changes in the animals exposed to a neutral novel context (not associated with US) as compared to 5US group (**Supplementary Figure** 1).

Overall, our data indicate that contextual fear extinction involves transient remodelling of the stOri neuronal circuit characterised by decreased density of dendritic spines with PSD-95 and upregulation of PSD-95 protein levels in the remaining dendritic spines. No significant synaptic changes were found in stRad, and only changes of PSD-95 in stLM.

### Synaptic changes in dCA1 during contextual fear extinction are homeostatic

Since we observed significant changes of dendritic spines and PSD-95 protein levels in dCA1 after fear extinction, in the following experiment we tested whether contextual fear extinction affected synaptic strength in dCA1 strata. To this end, field excitatory postsynaptic potentials (fEPSPs) were measured in acute hippocampal slices when the Shaffer collateral was stimulated by monotonically increasing stimuli (**Supplementary Figure** 2). The input-output curves showed no significant differences in the amplitude of fEPSP and fibre volley in stOri, stRad and stLM between the mice sacrificed immediately before or after fear extinction session (**Supplementary Figure** 2), indicating no global changes in synaptic strength. Thus our data indicate that extinction-induced remodelling of dendritic spines was homeostatic.

### Contextual fear extinction induces phosphorylation of PSD-95(S73) in dCA1

Phosphorylation of PSD-95(S73) has been associated with various forms of synaptic plasticity (Gardoni et al., 2006; Nowacka et al., 2020). To test whether contextual fear extinction induces phosphorylation of PSD-95(S73) in dCA1, we generated an antibody directed against this phosphorylation site (Figure 2A) (Gardoni et al., 2006). Mice underwent CFC and were sacrificed 24 hours later (5US), or after 15 or 30 minutes of the contextual fear extinction session (Ext15’ or Ext30’) (Figure 2B). The levels of PSD-95, phosphorylated PSD-95(S73) [phospho-PSD-95(S73)] and their colocalization were tested on the brain sections (Figure 2C). Total PSD-95, phospho-PSD-95(S73) and their colocalization levels were higher in the Ext15’, but not Ext30’, group as compared to the 5US animals (Figure D-F). Thus our data indicate that the alteration of PSD-95 protein levels during contextual fear extinction was accompanied by transiently increased phosphorylation of PSD-95(S73).

**Figure 2.**
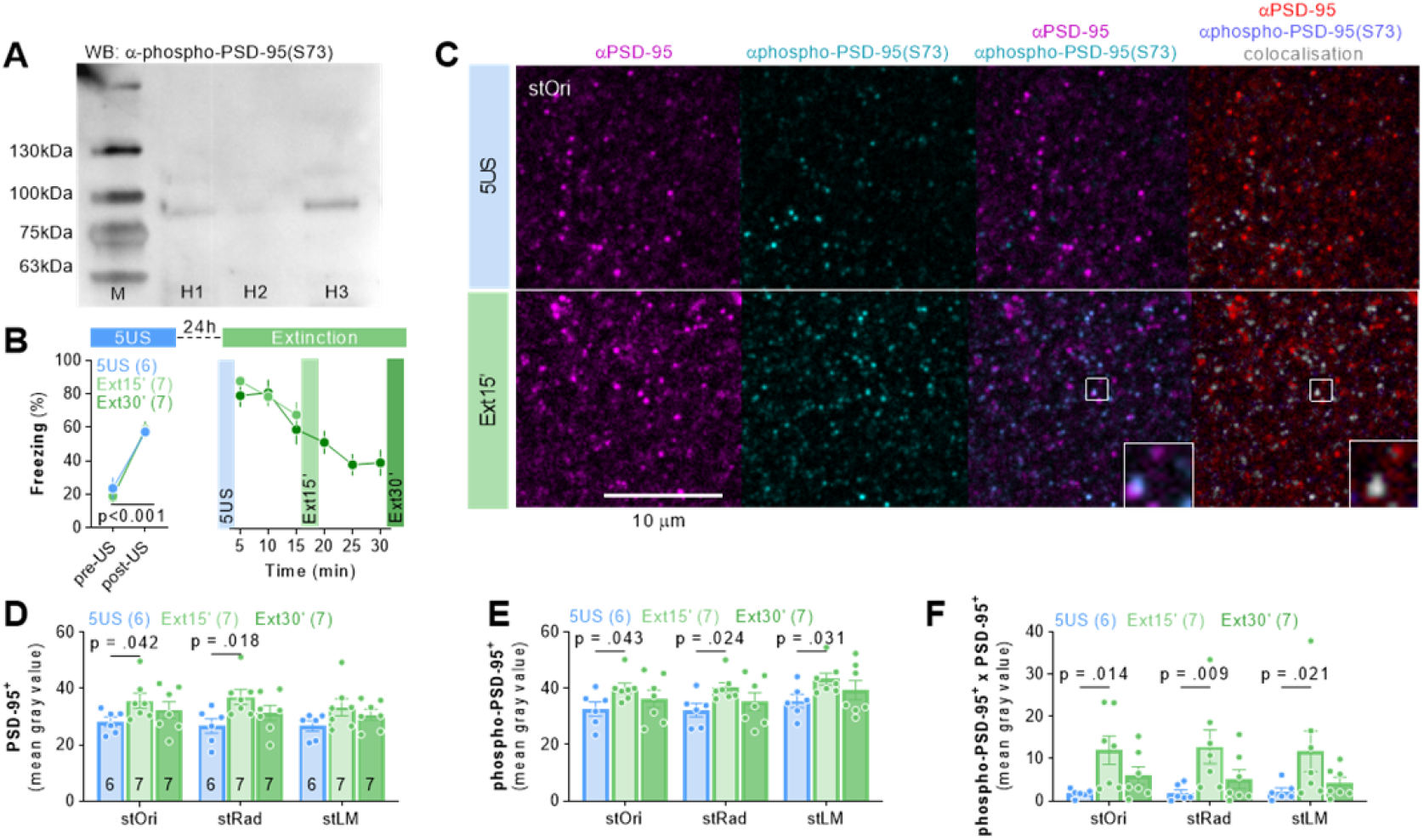
Contextual fear extinction induces transient phosphorylation of PSD-95(S73) in dCA1. **(A)** Western blot stained with phospho-PSD-95(S73)-specific antibody detects in the hippocampus homogenates (H1-3) proteins with approx. 95 kDA molecular weight. M, molecular weight marker. **(B)** Experimental timeline and freezing levels during training. Mice underwent CFC and were sacrificed 24 hours later (5US, n = 6) or after 15 or 30 minutes of a fear extinction session (Ext15’, n = 7; Ext30’, n = 7). **(C)** Representative confocal scans of the brain slices (stOri) immunostained with antibodies specific for PSD-95, phosphorylated PSD-95(S73) and their colocalization. **(D-F)** Quantification of the PSD-95 (two-way ANOVA, effect of training: F(2, 17) = 2.69, P = 0.097; effect of stratum: F(1,96, 33,3) = 3.83, P = 0.033), phospho-PSD-95(S73) (two-way ANOVA, effect of training: F(2, 17) = 2.20, P = 0.141; effect of stratum: F(1,24, 21,0) = 24.9, P < 0.001) and their colocalization levels (two-way ANOVA, effect of training: F(2, 17) = 4.08, P = 0.036; effect of stratum: F(2, 34) = 0.169, P = 0.845). Each dot represents one mouse. Means ± SEM are shown.

### PSD-95(S73) phosphorylation regulates PSD-95 protein levels during contextual fear extinction

To test whether phosphorylation of PSD-95(S73) regulates PSD-95 protein levels in dCA1 during fear extinction we used dCA1-targeted expression of phosphorylation-deficient PSD-95 with S73 mutated to alanine (S73A). We designed and produced adeno-associated viral vectors (AAV1/2) encoding wild-type PSD-95 protein under *Camk2a* promoter fused with mCherry (AAV1/2:CaMKII_PSD-95(WT):mCherry) (WT) or PSD-95(S73A) fused with mCherry (AAV1/2:CaMKII_PSD-95(S73A):mCherry) (S73A) (Nowacka et al., 2020) (**Supplementary Figure** 3). Mice underwent CFC (**Figure** 3A). The animals in all experimental groups showed increased freezing levels at the end of the training. Half of the mice were sacrificed 24 hours after CFC (5US). The remaining half were sacrificed after the 30-minut contextual fear extinction session (Ext). All animals showed high freezing levels at the beginning of the session, which decreased during the session. No effect of the virus was found (**Figure** 3A).

**Figure 3.**
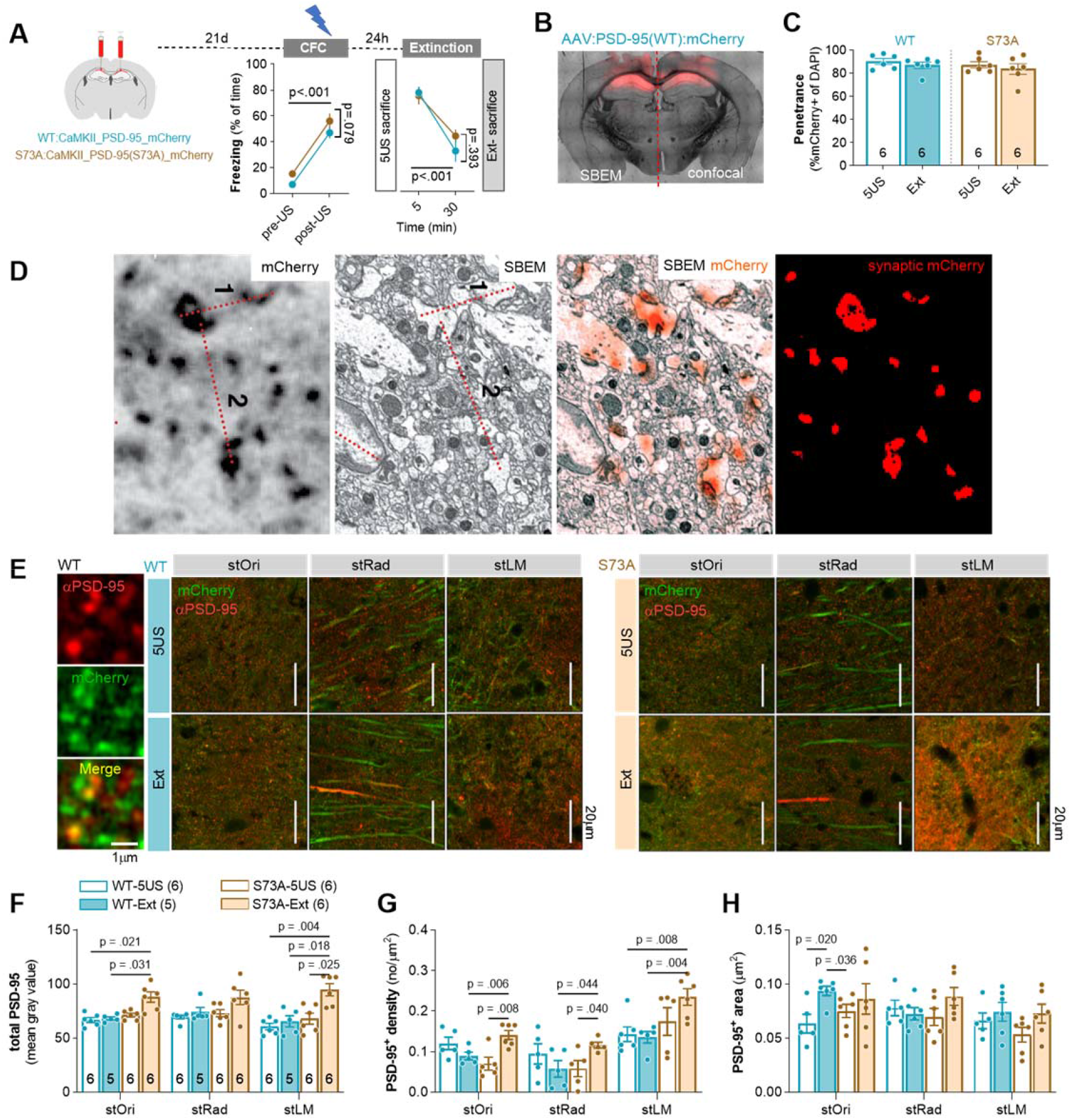
PSD-95(S73) is phosphorylated during fear extinction and this process is required for regulation of PSD-95 protein levels. **(A)** Experimental timeline and freezing during training. C57BL/6J male mice were stereotactically injected in the dCA1 with AAV1/2 encoding PSD-95(WT) (WT, n = 12) or PSD-95(S73A) (S73A, n = 12). Twenty one days later they underwent CFC (two-way repeated-measures ANOVA, effect of training: F(1, 30) = 269.4, P < 0.001, effect of virus: F(2, 30) = 2.815, P = 0.076) and were sacrificed 1 day after training (5US) or they were re-exposed to the training context without footshock and sacrificed (Ext) (two-way repeated-measures ANOVA, effect of training: F(1, 15) = 65.68, P < 0.001; effect of virus: F(2, 15) = 0.993, P = 0.393). **(B)** Microphotography of a brain with dCA1 PSD-95(WT):mCherry expression with illustration of the brain processing scheme**. (C)** Summary of data showing the viruses penetrance in dCA1 (sections used for confocal and SBEM analysis) (mice: 5US/Ext, WT = 6/5; S73A = 6/6). **(D)** Correlative confocal-electron microscopy analysis showing that exogenous PSD-95(WT) co-localises with PSDs. Single confocal scan of an exogenous PSD-95(WT) in dCA1, SBEM scan of the same area, superposition of confocal (orange) and SBEM images based on measured distances between large synapses (1 & 2), and thresholded synaptic PSD-95(WT) signal. Measurements: (confocal image) 1: 3.12 μm, 2: 4.97 μm; (SBEM image) 1: 2.98 μm, 2: 4.97 μm. **(E-H)** Analysis of PSD-95 expression after fear extinction training. **(E)** Representative confocal scans of the PSD-95 immunostaining. Means ± SEM are shown.

For each animal, half of the brain was chosen at random for confocal analysis of the PSD-95 protein levels, and the other half was processed for Serial Block-face Scanning Electron Microscopy (SBEM) (**Figure** 3B). The AAVs penetrance did not differ between the experimental groups (5US vs Ext) and reached over 80% of the cells in the analysed sections of dCA1 (**Figure** 3C). Correlative light and electron microscopy confirmed that the exogenous PSD-95 co-localised with postsynaptic densities (PSDs) representing a postsynaptic part of excitatory synapses, and only weak signal was present in dendrites (**Figure** 3D). We did not observe significant differences in total PSD-95 protein levels between the WT and S73A mice sacrificed before the fear extinction session. The total PSD-95 protein levels were not changed after fear extinction in the WT group, as compared to the WT mice sacrificed before the fear extinction session. However, PSD-95 levels were upregulated in all strata after the extinction session in the S73A mice, as compared to the WT Ext animals and the S73A 5US group (**Figure** 3F). As no significant differences in the mean PSD-95^+^ area were observed between the WT Ext and S73A Ext mice (**Figure** 3G), the differences in total PSD-95 levels likely resulted from higher density of PSD-95^+^ puncta in the S73A Ext group as compared to the S73A 5US and WT Ext animals (**Figure** 3F). Hence, exogenous PSD-95(S73A) protein impaired regulation of PSD-95^+^ density, indicating that phosphorylation of PSD-95(S73) controls PSD-95 levels during fear extinction.

### Phosphorylation of PSD-95(S73) regulates stOri synapses during fear extinction

To test whether phosphorylation of PSD-95(S73) regulates structural plasticity of excitatory synapses during contextual fear extinction we used SBEM. We reconstructed dendritic spines and PSDs in the stOri and determined dendritic spine density and volume as well as PSDs surface area [as a proxy of synaptic strength (Nusser et al., 1998; Noguchi et al., 2005; Katz et al., 2009)] and volume [as a proxy of the accumulated synaptic proteins (Borczyk et al., 2019)] (**Figure** 4A-D). In total, we reconstructed 159 spines from the brains of the WT mice sacrificed 24 hours after CFC (5US) (n=3), and 178 spines from the mice sacrificed after fear extinction (Ext) (n=3). For mice expressing S73A, 183 spines were reconstructed in the 5US group (n=3) and 160 Ext (n=3). Figure 4D shows reconstructions of dendritic spines from representative SBEM brick scans for each experimental group.

**Figure 4.**
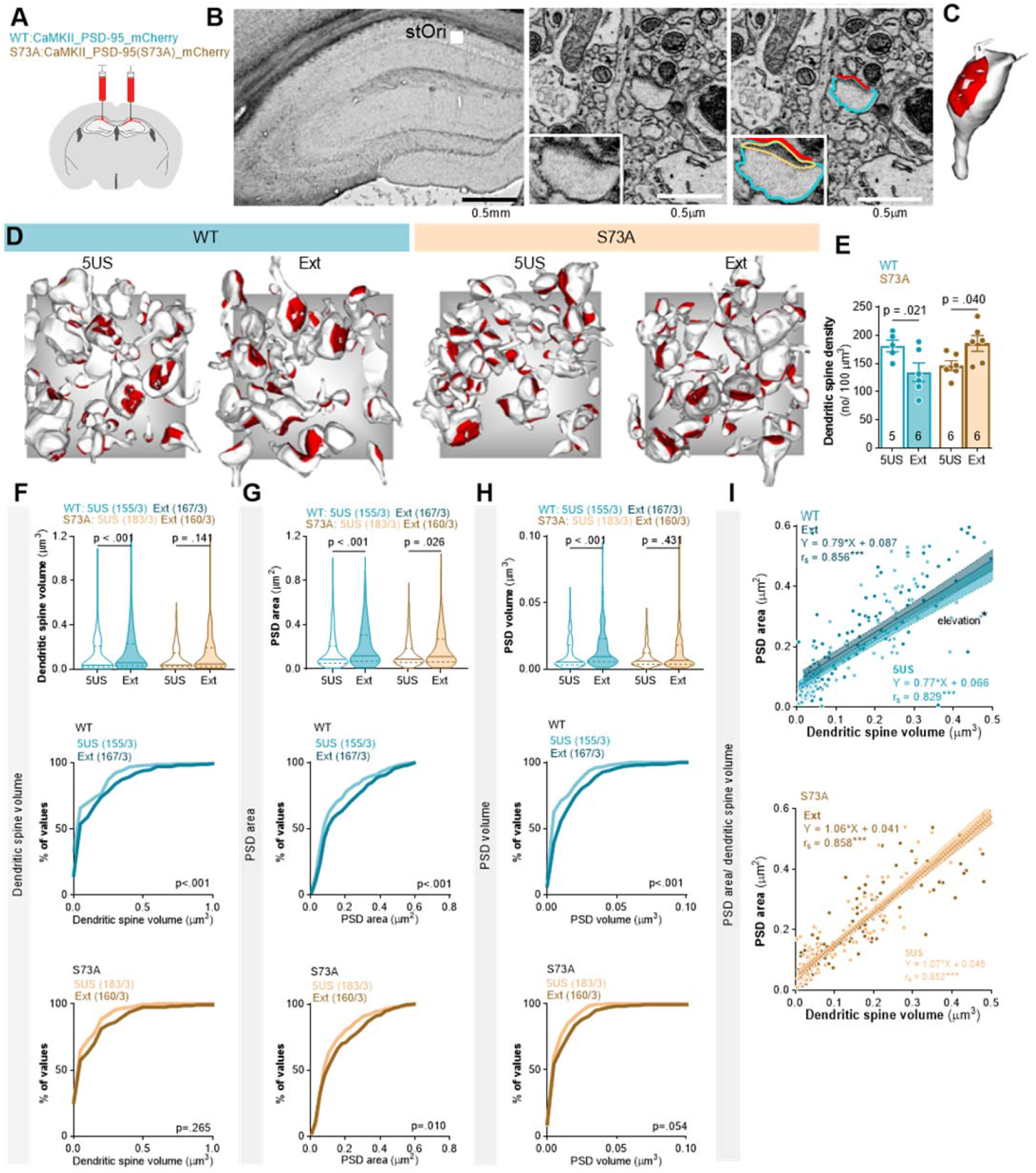
Phosphorylation of PSD-95(S73) regulates excitatory synapses during fear extinction. **(A)** Male mice were stereotactically injected in the dCA1 with AAV1/2 encoding PSD-95(WT) (WT, n = 12) or PSD-95(S73A) (S73A, n = 12). Twenty one days later they underwent CFC and were sacrificed 1 day after training (5US) or they were re-exposed to the training context for fear extinction (Ext). **(B-C)** The principles for SBEM analysis of the ultrastructure of dendritic spines and PSDs. **(B, left)** Microphotography of a dorsal hippocampus with the region of interest for analysis and tracing of a dendritic spine and PSD in stOri. **(B, right)** A representative trace of a dendritic spine (blue), PSD surface area (red) and volume (yellow), and **(C)** reconstruction of this dendritic spine. **(D)** Exemplary reconstructions of dendritic spines and their PSDs from SBEM scans in stOri. The grey background rectangles are x = 3 × y = 3 μm. Dendritic spines and PSDs were reconstructed and analysed in tissue bricks. **(E-I)** Summary of data showing: **(E)** mean density of dendritic spines (two-way ANOVA with LSD *post hoc* tests for planned comparisons, effect of training: F(1, 45) = 8.01, P = 0.007); **(F)** median dendritic spine volume (Mann-Whitney test, WT: U = 9766, P < 0.001; S73A: U = 13217, P = 0.141) and distributions of dendritic spine volumes (numbers of the analysed dendritic spines/mice are indicated) (Kolmogorov-Smirnov test, WT: D = 0.239, P < 0.001; S73A: D = 0.109, P = 0.265); **(G)** median PSD surface area (Mann-Whitney test, WT: U = 9948, P < 0.001; S73A: U = 46678, P = 0.024) and distributions of PSD surface areas (numbers of the analysed dendritic spines/mice are indicated) (Kolmogorov-Smirnov test, WT: D = 0.157, P < 0.001; S73A: D = 0.128, P = 0.010); **(H)** median PSD volume (Mann-Whitney test, WT: U = 9462, P < 0.001; S73A: U = 13621, P = 0.431) and distributions of PSD volumes (numbers of the analysed dendritic spines/mice are indicated)(Kolmogorov-Smirnov test, WT: D = 0.278, P < 0.001; S73A: D = 0.145, P = 0.054); **(I)** correlation of dendritic spine volume and PSD surface area (ANCOVA, WT: elevation, F(1, 319) = 4.256, P = 0.039; S73A: elevation, F(1, 340) = 0.603, P = 0.438; linear regression equations and Spearman correlation R are given for raw data). For E, each dot represents one tissue brick and means ± SEM are shown; for F, G, H (top) medians ± IQR are shown; for I, each dot represents an individual dendritic spine and regression lines ± 95% confidence intervals are shown.

Dendritic spine density was lower in the WT Ext group, as compared to the WT 5US mice (Figure 4E). Furthermore, the median values of dendritic spine volume, PSD surface area and PSD volume were higher after the extinction training in the WT group, as compared to the WT 5US mice. These changes were also indicated as shifts in the frequency distributions toward bigger values (Figure 4F-H, middle panels). We also observed the upward shift of the regression line describing the correlation between dendritic spine volume and PSD surface area in the WT Ext group, as compared to the WT 5US group (Figure 4I). Thus, in the WT group dendritic spines had relatively bigger PSDs after fear extinction than the dendritic spines of the same size in the 5US groups. Overall, the pattern of synaptic changes observed in the WT mice resembled the changes found in Thy1-GFP(M) animals after contextual fear extinction (Figure 1). The S73A mutation impaired fear extinction-induced downregulation of dendritic spine density as well as dendritic spine and PSD growth (Figure 4E-I). Altogether, our data indicate that PSD-95(S73) phosphorylation regulates both density and size of the excitatory synapses during contextual fear extinction.

### PSD-95(S73) phosphorylation in dCA1 is required for consolidation of contextual fear extinction memory

To test whether phosphorylation of PSD-95(S73) is necessary for consolidation of fear extinction memory, we used dCA1-targeted expression of S73A, WT or control AAV1/2 encoding mCherry under *Camk2a* promoter (Control). Two cohorts of mice with dCA1-targeted expression of the Control virus, WT or S73A, underwent CFC and fear extinction training. The first cohort underwent a short extinction training with one 30-minute extinction session (Ext) and 5-minute test of fear extinction memory (Test) (**Figure** 5A), while the second underwent an extensive fear extinction training with three 30-minute contextual fear extinction sessions on the days 2, 3, 4 (Ext1-3), followed by spontaneous fear recovery/ remote fear memory test on day 18, and further three extinction sessions on the days 18-20 (Ext4-6). Next, fear generalisation was tested in a context B (CtxB, day 22) (**Figure** 5D). The post-training analysis showed that the viruses were expressed in dCA1 (**Figure** 5I-J). The control virus was expressed in 85% of the dCA1 cells, WT in 88% and S73A in 87% (**Figure** 5J).

**Figure 5.**
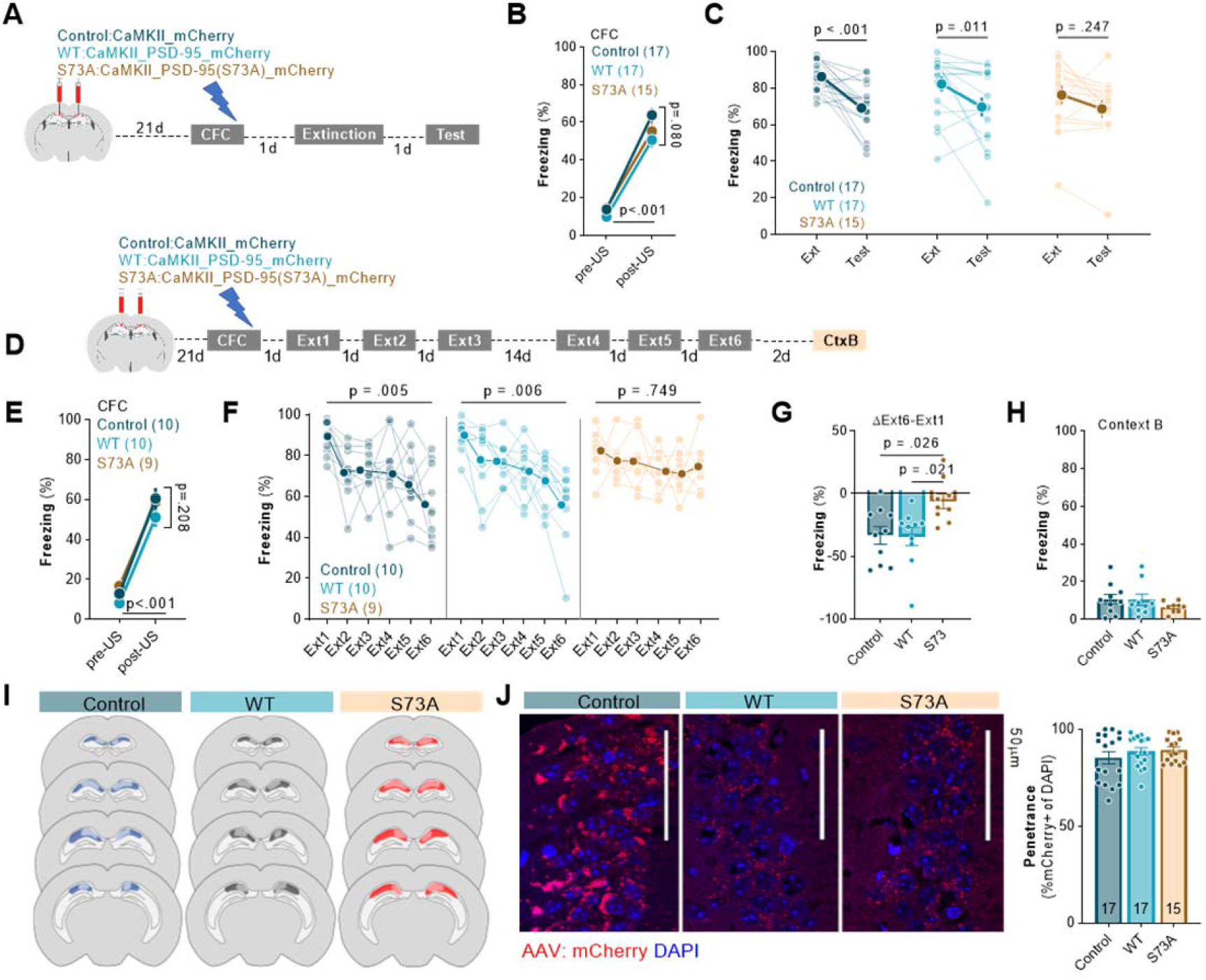
Phosphorylation of PSD-95(S73) in dCA1 is required for contextual fear extinction. **(A)** Experimental timeline of the short fear extinction training. C57BL/6J male mice were stereotactically injected in the dCA1 with AAV1/2 encoding mCherry (Control, n = 17), PSD-95(WT) (WT, n = 17) or PSD-95(S73A) (S73A, n = 15). Twenty one days after surgery mice underwent CFC. One day after CFC they were re-exposed to the training context in the absence of foot shock (Extinction). Consolidation of fear extinction memory was tested one day later in the same context (Test). **(B-C)** Summary of data showing percentage of freezing during **(B)** CFC, **(C)** Extinction and Test of the mice with dCA1-targeted expression of Control, WT or S73A (two-way repeated-measures ANOVA with Šídák’s multiple comparisons test, effect of time: F(1, 46) = 26.13, P < 0.001, genotype: F(2, 46) = 0.540, P = 0.586; time x genotype: F(2, 46) = 1.25, P = 0.296). **(D)** Experimental timeline of the extensive fear extinction training. Mice with dCA1-targeted expression of Control (n=10), WT (n=10) or S73A (n=9) underwent CFC, followed by six 30-min fear extinction sessions (Ext1-6) and one exposure to novel context without footshock (CtxB). **(E-H)** Summary of data showing freezing levels **(E)** during CFC, **(F)** after extensive fear extinction training (two-way repeated-measures ANOVA with Dunnett’s multiple comparisons test, effect of time: F(3.681, 95.70) = 13.01, P < 0.001; genotype: F(2, 26) = 1.23, P = 0.306; time x genotype: F(10, 130) = 1.49, P = 0.147), **(G)** the difference in freezing between Ext1 and Ext6 (one-way ANOVA with Tukey’s multiple comparisons test, F(2, 24.94) = 4.98, P = 0.016), and **(H)** during the test in the context B (Brown-Forsythe ANOVA test, F(2, 17.56) = 0.902, P = 0.428). **(I)** The extent of viral infection. **(J)** Single confocal scans of the stratum pyramidale of dCA1 of the mice expressing Control, WT and S73A and penetrance of the viruses. Means ± SEM are shown.

The analysis of the short extinction training (data pooled from two cohorts) showed that in all experimental groups freezing levels were low at the beginning of the training and increased after 5US delivery (**Figure** 5B). Furthermore, mice in all groups showed high freezing levels at the beginning of the Ext indicating similar levels of contextual fear memory acquisition. However, freezing measured during the Test was significantly decreased, as compared to the beginning of Ext, only in the Control and WT groups, not in the S73A animals (**Figure** 5C).

The analysis of freezing levels during the extensive fear extinction training showed high levels of freezing at the end of training and beginning of Ext1 for all experimental groups (**Figure** 5E-F). In the Control and WT groups, the freezing levels decreased over consecutive extinction sessions (Ext26) and were significantly lower as compared to Ext1, indicating formation of long-term fear extinction memory. We also found no spontaneous fear recovery after 14-day delay (Ext4 vs Ext3; Control, P = 0.806; WT, P = 0.248). In the S73A group, the extensive contextual fear extinction protocol did not reduce freezing levels measured at the beginning of Ext6 sessions, as compared to Ext1, indicating no fear extinction (**Figure** 5F). Accordingly we found significantly larger reduction of freezing after fear extinction training (ΔExt6-Ext1) in the controls and WT animals, as compared to the S73A group (**Figure** 5G). The freezing reaction was specific for the training context, as it was very low and similar for all experimental groups in the context B (**Figure** 5H). Thus, our data indicate that expression of the S73A in dCA1 does not affect fear memory formation, recall or generalisation but prevents contextual fear extinction even after extensive fear extinction training.

### αCaMKII autophosphorylation regulates contextual fear extinction and PSD-95 protein levels during contextual fear extinction

PSD-95(S73) is phosphorylated by αCaMKII (Gardoni et al., 2006; Steiner et al., 2008). To test the role of αCaMKII in PSD-95 protein regulation during fear extinction, autophosphorylation-deficient αCaMKII mutant mice (T286A) (Giese et al., 1998) and their wild-type (WT) littermates (males and females, in sex-balanced groups) were trained in CFC. They had similar and low levels of freezing in the novel context and freezing increased after 5US delivery (**Figure** 6A). Mice of both genotypes also showed high levels of freezing in the training context on the next day (Ext1), indicating contextual fear memory formation. However, when the mice were re-exposed to the training context for fear extinction (Ext2-3), the freezing levels of WT mice were significantly lower, as compared to Ext1, while the T286A mutants showed still high freezing. Thus, we confirmed that αCaMKII autophosphorylation is required for contextual fear memory extinction.

**Figure 6.**
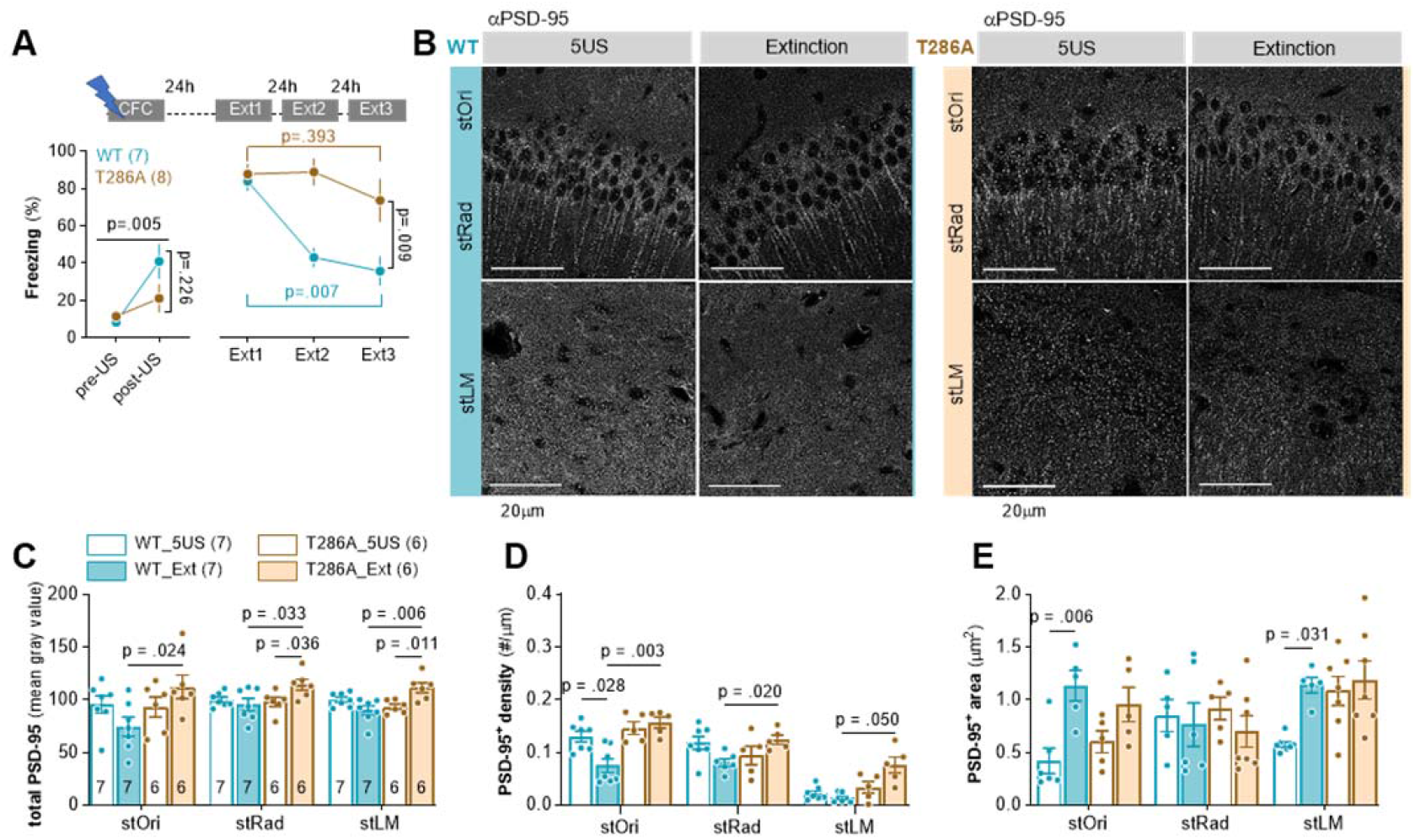
Autophosphorylation of αCaMKII is required for extinction of contextual fear and regulation of PSD-95 levels during fear extinction training. **(A)** Experimental timeline [WT and T286A underwent CTC and three fear extinction sessions (Ext1-3)] and percentage of freezing during CFC (two-way repeated-measures ANOVA, effect of time: F(1, 10) = 13.06, P = 0.005; effect of genotype: F(1, 10) = 1.66, P = 0.226) and Ext1-3 (WT/T286A = 7/8; sex-balanced groups) (two-way repeated-measures ANOVA with Šídák’s multiple comparisons test, effect of training: F(1,430, 18,59) = 14.96, P < 0.001; effect of genotype: F(1, 13) = 9.30, P = 0.009). **(B-E)** Analysis of PSD-95 expression in T286A mice. **(B)** Representative confocal scans of the brain slices immunostained for PSD-95. **(C-E)** Quantification of the PSD-95 protein levels: **(C)** total PSD-95 levels (three-way repeated-measures ANOVA with with Tukey’s *post hoc* test, effect of genotype x training: F(1, 22) = 15.03, P < 0.001); **(D)** density of PSD-95^+^ puncta (three-way repeated-measures ANOVA with Tukey’s *post hoc* test, effect of genotype x training: F(1, 59) = 23.43, P < 0.001); **(E)** area of PSD-95^+^ puncta (three-way repeated-measures ANOVA with Tukey’s *post hoc* test, effect of region x training: F(2, 35) = 6.858, P = 0.003). Mice: 5US/Ext, WT= 7/7; T286A = 6/6. Means ± SEM are shown.

Next, a second cohort of WT and T286A mice was trained and the animals were sacrificed 24 hours after training (5US) or after fear extinction session (Ext). The total levels of PSD-95 were not affected by the fear extinction session in WT mice (**Figure** 6C). However, PSD-95 levels were higher in all strata of dCA1 in the T286A Extinction group, as compared to WT animals sacrificed after extinction, and T286A mutants sacrificed before extinction. The density of PSD-95^+^ puncta was decreased, while mean area of the puncta increased in stOri in the WT Ext animals as compared to the WT 5US mice (**Figure** 6D-E). Moreover, in the T286A Ext group the density of PSD-95^+^ puncta was higher in all strata of dCA1, as compared to the WT Ext animals (**Figure** 6D). Thus, this experiment supports the hypothesis that αCaMKII autophosphorylation is required for extinction-induced regulation of PSD-95^+^ density.

## DISCUSSION

We have investigated the role of dCA1 PSD-95(S73) phosphorylation in contextual fear extinction. Our study showed that: (1) contextual fear extinction induces transient changes of dCA1 PSD-95 protein levels and dendritic spines in a stratum-specific manner. The most pronounced changes are observed in stOri; (2) contextual fear extinction induces phosphorylation of PSD-95(S73) in all dCA1 strata; (3) Expression of the exogenous, phosphorylation-deficient PSD-95(S73A) in dCA1 deregulates PSD-95 protein levels and synaptic remodelling induced by extinction of fear memories; (4) dCA1 PSD-95(S73A) impaires long-term, contextual fear extinction memory, but not for fear memory formation or recall; (5) Phosphorylation-deficient αCaMKII(T286A) impairs contextual fear extinction and regulation of dCA1 PSD-95 protein levels during fear extinction.

Here, we demonstrate that contextual fear extinction transiently increases phospho-PSD-95(S73) levels and induces rapid downregulation of the synapses with PSD-95 as well as growth of the remaining synapses in stOri. These synaptic processes are homeostatic - without the changes of total PSD-95 levels and synaptic strength. Such synaptic plasticity alludes to the Hebbian strengthening of activated synapses and heterosynaptic weakening of adjacent synapses (Royer and Paré, 2003; El-Boustani et al., 2018). Our study is the first demonstration of the homeostatic plasticity of dendritic spines during attenuation of fear memories. We also show that extinction-induced downregulation of stOri synapses, as well as regulation of PSD-95 protein levels, are impaired by the expression of phosphorylation-deficient PSD-95(S73A). These observations indicate that phosphorylation of PSD-95(S73) is a key step in the regulation of the dCA1 circuit during fear extinction. There are several important limitations of our study. Firstly, using phospho-S73 antibody we cannot exclude that other MAGUKs are detected (due to the similar LERGN**S**GLGFS sequence). However, the role of phospho-PSD-95(S73) in contextual fear extinction is supported by the fact that there is increased colocalization of PSD-95 and phospho-S73 during extinction. Moreover, using *ex vivo* analyses we cannot unequivocally indicate whether PSD-95(S73A) prevents elimination of dendritic spines and PSD-95 proteins, or changes the balance of the synapses by enhancing synaptogenesis and protein synthesis. We believe, however, that the first scenario is more likely and this conclusion is supported by several observations. Firstly, PSD-95(S73A) does not affect synaptic strengthening (Steiner et al., 2008), but PSD-95(S73) phosphorylation allows for dissociation of PSD-95 from the complex with GluN2A, destabilisation of PSD and termination of synaptic growth after NMDAR stimulation (Gardoni et al., 2006; Steiner et al., 2008) as well as downregulation of PSD-95 levels during NMDAR-LTD (Nowacka et al., 2020). Secondly, both dCA1 phosphorylation of PSD-95(S73) and protein degradation, but not protein synthesis, are necessary for contextual fear extinction (Fischer, 2004; Lee et al., 2008).

Our experiments are the first to show that phosphorylation of PSD-95(S73) in dCA1 is required for extinction of contextual fear memories. Strikingly, the contextual fear memory cannot be updated even when the animals with dCA1 PSD-95(S73A) mutation undergo six 30-minute extinction sessions. We also show that dCA1 PSD-95(S73A) does not affect mice activity, long-term fear memory formation and recall, context-independent fear generalisation or fear recovery after 14-day delay, pointing towards engagement of PSD-95(S73) phosphorylation only during extinction of contextual fear. This conclusion seemingly contradicts the study demonstrating that ligand binding-deficient PSD-95 knockin mice have enhanced contextual fear memory formation and impaired longterm memory retention (Nagura et al., 2012; Fitzgerald et al., 2015). However, even though the behavioural phenotype of PSD-95 KI mice was supported by LTP analysis in dCA1 (Nagura et al., 2012; Fitzgerald et al., 2015), it is unknown whether the mouse phenotype relies on the CA1 plasticity as the mutation was global. Furthermore, it is possible that PSD-95 KI and PSD-95(S73A) impact different stages of contextual fear memory. In agreement with our findings, the signalling pathways downstream of NMDAR-PSD-95 complex in the dorsal CA3 and DG regulate contextual fear extinction (Li et al., 2017; Cai et al., 2018). In particular, translocation of PSD-95 from NMDAR to TrkB, and increased PSD-95-TrkB interactions, promote extinction, while competing NMDAR-PSD-95-nNOS interactions hinder contextual fear extinction (Cai et al., 2018). Since PSD-95(S73A) mutation prolongs NMDAR-PSD-95 interactions (Gardoni et al., 2006) it may limit interactions of PSD-95 with TrkB and fear extinction. To support this hypothesis we also show that autophosphorylation of αCaMKII, the key enzyme activated by NMDAR, is required for extinction-induced regulation of PSD-95 levels and fear extinction.

Our data show that the extinction of contextual fear affects PSD-95 protein levels and dendritic spines predominantly in the stOri. This indicates that the extinction-induced synaptic remodelling is strikingly different from the changes observed immediately after contextual fear memory encoding where transient synaptogenesis is observed in the stRad (Radwanska et al., 2011). These observations support the idea that different CA1 inputs are involved in memory formation and extinction. CA3 neurons project to the stRad and stOri regions of CA1 pyramidal neurons, the nucleus reuniens (Re) projects to the stOri and stLM, and the entorhinal cortex (EC) projects to the stLM (Ishizuka et al., 1990; Kajiwara et al., 2008; Hoover and Vertes, 2012; Vertes et al., 2015). Thus, the pattern of synaptic changes induced by contextual fear extinction co-localises with the domains innervated by the Re and EC, suggesting that these inputs are regulated during contextual fear extinction. In agreement with our observations, previous data showed that the EC is activated during and required for contextual fear extinction in animal models (Bevilaqua et al., 2006; Baldi and Bucherelli, 2015). Human studies also showed that EC-CA1 projections are activated by cognitive prediction error (that may drive memory extinction), while CA3-CA1 projections are activated by memory recall without prediction errors (Bein et al., 2020). The role of the Re in fear memory encoding, retrieval, extinction and generalisation has been demonstrated (Xu and Sudhof, 2013; Ramanathan et al., 2018; Troyner and Bertoglio, 2021). Still, it has to be established whether the plasticity of dCA1 synapses is specific to Re and/or EC projections.

Our findings add up to the previous studies investigating the molecular processes in dCA1 that are specific and required for contextual fear extinction, but not for fear memory consolidation, including regulation of ERK, CB1, and CBEP (Berger-Sweeney et al., 2006; Bitencourt et al., 2008; de Oliveira Alvares et al., 2008; Pamplona et al., 2008; Tronson et al., 2009; Radulovic and Tronson, 2010). Interestingly, other processes, such as protein synthesis and c-Fos expression, are necessary for contextual fear consolidation and reconsolidation, but not extinction (Fischer, 2004; Lattal and Abel, 2004; Mamiya et al., 2009; Tronson et al., 2009). Thus, although it is not surprising that distinct molecular cascades and cell circuits contribute to fear memory formation/recall and extinction (Tronson et al., 2009; Lacagnina et al., 2019), it remains puzzling how synaptic plasticity, without concomitant translation, contributes to contextual fear extinction. This observation points towards the role of protein synthesis-independent short-term plasticity, or protein degradation (Lee et al., 2008), in contextual fear extinction memory. The role of short-term plasticity in contextual fear extinction is supported by the observations that PSD-95(S73) phosphorylation and synaptic remodelling induced by fear extinction are transient. Similar short plasticity was observed by other groups upon recall of drug-paired memories (Gipson et al., 2013a, 2013b). Still it has to be clarified in the future studies how short-term dCA1 plasticity can support long-term fear extinction memory.

## Conclusions

Our study demonstrates that extinction of contextual fear memories relies on rapid and transient synaptic plasticity in dCA1 that requires PSD-95(S73) phosphorylation. Thus our study supports the hypothesis that NMDAR-dependent plasticity in dCA1 is required to detect and resolve contradictory or ambiguous memories when spatial information is involved (Bannerman et al., 2014), the comparator view of hippocampal function (Gray, 1982; Grossberg and Merrill, 1992) as well as the observations that the hippocampus processes surprising events and prediction errors (Ploghaus et al., 2000; Kumaran and Maguire, 2006; Huh et al., 2009; Bein et al., 2020). Since new or long-lasting memories may be repeatedly reorganised upon recall (Nader et al., 2000; Schafe et al., 2001), the molecular and cellular mechanisms involved in extinction of the existing fearful memories provide excellent targets for fear memory impairment therapies. In particular, understanding the mechanisms that underlie contextual fear extinction may be relevant for post-traumatic stress disorder treatment.

## Acknowledgments, Funding and Disclosure

This work was supported by a National Science Centre (Poland) Grant No. 2015/19/B/NZ4/02996 and 2020/38/A/NZ4/00483 to KR. PRELUDIUM Grant No. 2016/21/N/NZ4/03304 to MZ and PRELUDIUM Grant No. 2015/19/N/NZ4/03611 to KŁ. TW was supported by the National Science Centre (Poland) (Grant No. 2017/26/E/NZ4/00637). The project was carried out using CePT infrastructure financed by the European Union - The European Regional Development Fund within the Operational Program “Innovative economy” for 2007-2013.

MZ, MB, KFT and KR designed the experiments; MZ, MB, AC, MNS, AN, MŚ, KŁ, KFT, TW and AS performed the experiments; MZ, MB, MŚ, KŁ, ES, KFT, JW, TB and KR analysed data. MZ, MB and KR drafted the manuscript. All authors had critical input to the final version of the manuscript. Authors report no financial interests or conflicts of interest. Light and electron microscopy experiments were performed at the Laboratory of Imaging Tissue Structure and Functions, Nencki Institute.

## MATERIALS AND METHODS

### Animals

C57BL/6J male mice were purchased from Białystok University, Poland. Thy1-GFP(M) (The Jackson Laboratory, JAX:007788, RRID:IMSR_JAX:007788) mutant mice were bred as heterozygotes at Nencki Institute, and PCR genotyped as previously described (Feng et al., 2000). αCaMKII-T286A mutant mice were bred as heterozygotes at Nencki Institute, and PCR genotyped as previously described (Giese et al., 1998). All mice in the experiments were 10-week old at the beginning of the experiments. The mice were housed in groups of two to six and maintained on a 12 h light/dark cycle with food and water *ad libitum*. All experiments with transgenic mice used approximately equal numbers of males and females. The experiments were undertaken according to the Animal Protection Act of Poland and approved by the I Local Ethics Committee (261/2012 and 829/2019 Warsaw, Poland).

### Contextual fear conditioning (CFC)

The animals were trained in a conditioning chamber (Med Associates Inc, St Albans, USA) in a soundproof box. The chamber floor had a stainless steel grid for shock delivery. Before training, the chamber was cleaned with 70% ethanol, and a paper towel soaked in ethanol was placed under the grid floor. To camouflage background noise in the behavioural room, a white noise generator was placed inside the soundproof box.

On the conditioning day, the mice were brought from the housing room into a holding room to acclimatise for 30 min before training. Next, mice were placed in the training chamber, and after a 148 s introductory period, a foot shock (2 s, 0.7 mA) was presented. The shock was repeated 5 times, at 90 s inter-trial intervals. Thirty seconds after the last shock, the mouse was returned to its home cage. Contextual fear memory was tested and extinguished 24 h after training by re-exposing mice to the conditioning chamber for 30 minutes without US presentation, followed by the second 5-minute test session on the following day. During extensive contextual fear extinction, 30-minute fear extinction sessions were repeated on days 2, 3, 14, 15, and 16. Moreover mice activity and freezing were tested in context B (Ctx B) on day 17. A video camera was fixed inside the door of the sound attenuating box for the behavior to be recorded and scored. Freezing behavior (defined as complete lack of movement, except respiration) and locomotor activity of mice were automatically scored. The experimenters were blind to the experimental groups.

### Stereotactic surgery

Mice were fixed in a stereotactic frame (51503, Stoelting, Wood Dale, IL, USA) and kept under isoflurane anesthesia (5% for induction, 1.5-2.0% during surgery). Adeno-associated viruses, serotype 1 and 2, (AAV1/2), solutions were injected into the dorsal CA1 area (Paxinos & Franklin 2001) at coordinates in relation to Bregma (AP, −2.1mm; ML, ±1.1 mm; DV, −1.3mm). 450 nl of AAV solutions were injected into the CA1 through a beveled 26 gauge metal needle, and 10 μl microsyringe (SGE010RNS, WPI, USA) connected to a pump (UMP3, WPI, Sarasota, USA), and its controller (Micro4, WPI, Sarasota, USA) at a rate 50 nl/ min. The needle was then left in place for 5 min, retracted +100 nm DV, and left for an additional 5 min to prevent unwanted spread of the AAV solution. Titers of AAV1/2 were: αCaMKII_PSD-95(WT):mCherry (PSD-95(WT)): 1.35 x10^9^/μl, αCaMKII_PSD-95(S73A):mCherry (PSD-95(S73A)): 9.12 x10^9^/μl), αCaMKII_mCherry (mCherry): viral titer 7.5 x10^7^/μl (obtained from Karl Deisseroth’s Lab). Mice were allowed to recover from anaesthesia for 2-3 h on a heating pad and then transferred to individual cages where they stayed until complete skin healing, and next, they were returned to the home cages. The viruses were prepared at the Nencki Institute core facility, Laboratory of Animal Models. After training, the animals were perfused with 4% PFA in PBS and bain sections from the dorsal hippocampus were immunostained for PSD-95 and imaged with Zeiss Spinning Disc confocal microscope (magnification: 10x) to assess the extent of the viral expression and PSD-95 expression.

### Immunostaining

Mice were anaesthetised and perfused with cold phosphate buffer pH 7.4, followed by 0.5% 4% PFA in phosphate buffer. Brains were removed and postfixed o/n in 4°C. Brains were kept in 30% sucrose in PBS for 72h. Coronal brain sections were prepared using cryosectioning (40 μm thick, Cryostat CM1950, Leica Biosystems Nussloch GmbH, Wetzlar, Germany) and stored in a cryoprotecting solution in −20°C (PBS, 15% sucrose (Sigma-Aldrich), 30% ethylene glycol (Sigma-Aldrich), and 0.05% NaN_3_ (SigmaAldrich). Before staining, sections were washed 3 × PBS and blocked for 1 hour at room temperature (RT) in 5% NDS with 0.3% Triton X-100 in PBS and then incubated o/n, 4°C with PSD-95 primary antibodies (1:500, Millipore, MAB1598, RRID:AB_11212185) and/or rabbit anti-mCherry primary antibodies (1:500, Abcam, ab167453, RRID:AB_2571870) and/or rabbit P-Ser73_PSD-95 primary antibodies (1:12, Davids Biotechnology, A061). On the second day slices were washed 3 × PBS with 0,3% Trition X-100 and incubated for 90 minutes with secondary antibodies conjugated with anti-mouse AlexaFluor 555 (1:500, Invitrogen, A31570, RRID:AB_2536180) and/or anti-rabbit AlexaFluor 555 (1:500, Invitrogen, A31572, RRID:AB_162543) and/or anti-rabbit Alexa Fluor 647 (1:500, Invitrogen, A31573, RRID:AB_2536183). Slices were then mounted on microscope slides (Thermo Fisher Scientific) and covered with coverslips in Fluoromount-G medium with DAPI (00-4959-52, Invitrogen).

### Phospho-PSD-95(S73)-specific antibody

Phospho-epitope-specific serum against phosphorylated PSD-95(S73) was raised in a rabbit using the synthetic phosphopeptide LERGN(Sp)GLGFS. The antibody was prepared and affinity-purified by Davids Biotechnologie (Regensburg, Germany).

### Confocal microscopy and image quantification

The microphotographs of dendritic spines in the Thy1-GFP(M) mice, fluorescent PSD-95 and phospho-PSD-95(S73) immunostaining were taken on a Spinning Disc confocal microscope (63 × oil objective, NA 1.4, pixel size 0.13 μm × 0.13 μm) (Zeiss, Göttingen, Germany). We took microphotographs (16 bit, z-stacks of 12-48 scans; 260 nm z-steps) of 6 dendrites per region per animal from stratum oriens (stOri), stratum radiatum (stRad) and stratum lacunosum-moleculare (stLM) (in the middle of the strata) of dCA1 pyramidal neurons (AP, Bregma from −1.7 to 2.06). The PSD-95 fluorescent immunostaining after AAV overexpression was analysed with Zeiss LSM 800 microscope equipped with Airy-Scan detection (63× oil objective and NA 1.4, pixel size 0.13 μm × 0.13 μm, 8 bit) (Zeiss, Göttingen, Germany). A series of 18 continuous optical sections (67.72 μm × 67.72 μm), at 0.26 μm intervals, were scanned along the z-axis of the tissue section. Six to eight z-stacks of microphotographs were taken per animal per region, from every sixth section through dCA1. Each dendritic spine was manually outlined, and the spine area was measured with ImageJ 1.52n software measure tool. Custom-written Python scripts were used to analyse the mean grey value of PSD-95^+^ puncta per dendritic spine, total PSD-95 levels (as an image mean gray value), and PSD-95^+^ puncta density and size.

### Serial Face-block Scanning Electron Microscopy (SBEM)

Mice were transcardially perfused with cold phosphate buffer pH 7.4, followed by 0.5% EM-grade glutaraldehyde (G5882 Sigma-Aldrich) with 2% PFA in phosphate buffer pH 7.4 and postfixed overnight in the same solution. Brains were then taken out of the fixative and cut on a vibratome (Leica VT 1200) into 100 μm slices. Slices were kept in phosphate buffer pH 7.4, with 0.1% sodium azide in 4°C. For AAV-injected animals, the fluorescence of exogenous proteins was confirmed in all slices by fluorescent imaging. Then, slices were washed 3 times in cold phosphate buffer and postfixed with a solution of 2% osmium tetroxide (#75632 Sigma-Aldrich) and 1.5 % potassium ferrocyanide (P3289 Sigma-Aldrich) in 0.1 M phosphate buffer pH 7.4 for 60 min on ice. Next, samples were rinsed 5 × 3 min with double distilled water (ddH_2_O) and subsequently exposed to 1% aqueous thiocarbohydrazide (TCH) (#88535 Sigma) solution for 20 min. Samples were then washed 5 × 3 min with ddH_2_O and stained with osmium tetroxide (1% osmium tetroxide in ddH_2_O, without ferrocyanide) for 30 min in RT. Afterward, slices were rinsed 5 × 3 min with ddH_2_O and incubated in 1% aqueous solution of uranium acetate overnight in 4°C. The next day, slices were rinsed 5 × 3 min with ddH_2_O, incubated with lead aspartate solution (prepared by dissolving lead nitrate in L-aspartic acid as previously described (Deerinck et al., 2010)) for 30 min in 60°C and then washed 5 × 3 min with ddH_2_O and dehydration was performed using graded dilutions of ice-cold ethanol (30%, 50%, 70%, 80%, 90%, and 2 × 100% ethanol, 5 min each). Then slices were infiltrated with Durcupan resin. A(17 g), B(17 g) and D(0,51 g) components of Durcupan (#44610 Sigma-Aldrich) were first mixed on a magnetic stirrer for 30 min and then 8 drops of DMP-30 accelerator (#45348 Sigma) were added (Knott et al., 2009). Part of the resin was then mixed 1:1 (v/v) with 100% ethanol and slices were incubated in this 50% resin on a clock-like stirrer for 30 min in RT. The resin was then replaced with 100% Durcupan for 1 hour in RT and then 100% Durcupan infiltration was performed o/n with constant slow mixing. The next day, samples were infiltrated with freshly prepared resin (as described above) for another 2 hours in RT, and then embedded between flat Aclar sheets (Ted Pella #10501-10). Samples were put in a laboratory oven for at least 48 hours at 65°C for the resin to polymerize. After the resin hardened, the Aclar layers were separated from the resin embedded samples, dCA1 region was cut out with a razorblade. Caution was taken for the piece to contain minimal resin. Squares of approximately 1 × 1 × 1 mm were attached to aluminium pins (Gatan metal rivets, Oxford instruments) with very little amount of cyanacrylamide glue. After the glue dried, samples were mounted to the ultramicrotome to cut 1 μm thick slices. Slices were transferred on a microscope slide, briefly stained with 1% toluidine blue in 5% borate and observed under a light microscope to confirm the region of interest (ROI). Next, samples were grounded with silver paint (Ted Pella, 16062-15) and pinned for drying for 4 – 12 hours, before the specimens were mounted into the 3View2 chamber.

SBEM *imaging and 3D reconstructions*. Samples were imaged with Zeiss SigmaVP (Zeiss, Oberkochen, Germany) scanning electron microscope equipped with 3View2 chamber using a backscatter electron detector. Scans were taken in the middle portion of dCA1 stOri. From each sample, 200 sections were collected (thickness 60 nm). Imaging settings: high vacuum with EHT 2.9-3.8 kV, aperture: 20 μm, pixel dwell time: 3 μs, pixel size: 5 – 6.2 nm. Scans were aligned using the ImageJ software (ImageJ -> Plugins -> Registration -> StackReg) and saved as .tiff image sequence. Next, aligned scans were imported to Reconstruct software (Fiala 2005), available at http://synapses.clm.utexas.edu/tools/reconstruct/reconstruct.stm (Synapse Web Reconstruct, RRID:SCR_002716). Dendritic spine density was analysed from 3 bricks per animal with the unbiased brick method (Fiala and Harris 2001) per tissue volume. Brick dimensions 3 × 3 × 3 μm were chosen to exceed the length of the largest profiles in the data sets at least twice. To calculate the density of dendritic spines, the total volume of large tissue discontinuities was subtracted from the volume of the brick. The density of dendritic spines was normalised to AAV1/2 penetrance.

A structure was considered to be a dendritic spine when it was a definite protrusion from the dendrite, with electron-dense material (representing postsynaptic part of the synapse, PSD) on the part of the membrane that opposed an axonal bouton with at least 3 vesicles within a 50-nm distance from the cellular membrane facing the spine. For 3D reconstructions, PSDs and dendritic spines in one brick were reconstructed for each sample. PSDs were first reconstructed and second, their dendritic spines were outlined. To separate dendritic spine necks from the dendrites, a cut-off plane was used approximating where the dendritic surface would be without the dendritic spine. PSD volume was measured by outlining dark, electron-dense area on each PSD containing section (Borczyk et al., 2019). The PSD area was measured manually according to the Reconstruct manual. All non-synaptic protrusions were omitted in this analysis. For multi-synaptic spines, the PSD areas and volumes were summed.

### Correlative light-electron microscopy (CLEM)

CLEM workflow was based on a previously established protocol with some modifications (Bishop et al., 2011). Mice infused with PSD-95(WT) in the CA1 were perfused as described above. Brains were then removed and postfixed o/n in 4°C. 100 μm thick brain slices were cut on a vibratome and embedded in low melting point agarose in phosphate buffer and mounted into imaging chambers. mCherry fluorescence in the stRad was photographed using Zeiss LSM800, z-stacks of 60 images (60 μm thick) at 63 × magnification. Next, the slice was transferred under the 2P microscope (Zeiss MP PA Setup), where a Chameleon laser was used to brand mark the ROI (laser length 870 nm, laser power 85%, 250 scans of each line). Then, SBEM staining was performed as described above. The resin-embedded hippocampus was then divided into 4 rectangles and each was mounted onto metal pins to locate the laser-induced marks. SBEM scanned within the laser marked frame. The fluorescent image was overlaid onto the SBEM image using dendrites and cell nuclei as landmarks using ImageJ 1.48k software (RRID:SCR_003070).

### Electrophysiology

Mice were deeply anaesthetised with Isoflurane, decapitated and the brains were rapidly dissected and transferred into ice-cold cutting artificial cerebrospinal fluid (ACSF) consisting of (in mM): 87 NaCl, 2.5 KCl, 1.25 NaH_2_PO_4_, 25 NaHCO_3_, 0.5 CaCl_2_, 7 MgSO_4_, 20 D-glucose, 75 sacharose equilibrated with carbogen (5% CO_2_/95% O_2_). The brain was cut to two hemispheres and 350 μm thick coronal brain slices were cut in ice-cold cutting ACSF with Leica VT1000S vibratome. Slices were then incubated for 15 min in cutting ACSF at 32°C. Next the slices were transferred to recording ACSF containing (in mM): 125 NaCl, 2.5 KCl, 1.25 NaH_2_PO_4_, 25 NaHCO_3_, 2.5 CaCl_2_, 1.5 MgSO_4_, 20 D-glucose equilibrated with carbogen and incubated for minimum 1 hour at room temperature (RT).

Extracellular field potential recordings were recorded in a submerged chamber perfused with recording ACSF in RT. The synaptic potentials were evoked with a Stimulus Isolator (A.M.P.I Isoflex) with a concentric bipolar electrode (FHC, CBARC75) placed in the stOri of CA2 on the experiment. The stimulating pulses were delivered at 0.1 Hz and the pulse duration was 0.3 ms. Recording electrodes (resistance 1-4 MΩ) were pulled from borosilicate glass (WPI, 1B120F-4) with a micropipette puller (Sutter Instruments, P-1000) and filled with recording ACSF. The recording electrodes were placed in stOri of dCA1. Simultaneously, a second recording electrode was placed in the stratum pyramidale to measure population spikes. For each slice, the recordings were done in stOri. Recordings were acquired with MultiClamp 700B amplifier (Molecular Devices, California, USA), digitised with Digidata 1550B (Molecular Devices, California, USA) and pClamp 10.7 Clampex 10.0 software (Molecular Devices, California, USA). Input/output curves were obtained by increasing stimulation intensity by 25 μA in the range of 0-300 μA. All electrophysiological data were analysed with AxoGraph 1.7.4 software (Axon Instruments, U.S.A). The amplitude of fEPSP, relative amplitude of population spikes and fibre volley were measured.

### Statistics

Data are presented as mean ± standard error of the mean (SEM) for populations with normal distribution or as median ± interquartile range (IQR) for populations with non-normal distribution. An animal was used as a biological replication in all experiments except for the dendritic spine size distribution analysis. When the data met the assumptions of parametric statistical tests, results were analysed by one-or repeated measures two-way ANOVA, followed by Tukey’s or Fisher’s *post hoc* tests, where applicable. Data were tested for normality by using the Shapiro-Wilk test of normality and for homogeneity of variances by using the Levene’s test. For repeated-measure data with missing observation, a linear mixed model was used to analyse the results, followed by pairwise comparisons with Sidak adjustment for multiple comparisons. Areas of dendritic spines and PSDs did not follow normal distributions and were analysed with the Kruskal-Wallis test. Frequency distributions of PSD area to the spine volume ratio were compared with the Kolmogorov-Smirnov test. Correlations were analysed using Spearman correlation (Spearman r (s_r_) is shown), and the difference between slopes or elevation between linear regression lines was calculated with ANCOVA. Differences between the experimental groups were considered statistically significant if P < 0.05. Analyses were performed using the Graphpad Prism 9. Mice were excluded from the analysis only if they did not express the tested virus in the target region.

| Key Resources Table |  |  |  |  |
| --- | --- | --- | --- | --- |
| Reagent type (species) or resource | Designation | Source or reference | Identifiers | Additional information |
| strain, strain background (*Mus musculus*, male) | Thy1-GFP(M) | PMID: 11086982 |  |  |
| strain, strain background (*Mus musculus*, male) | C57BL/6J | Bialystok University, Poland | IMSR_JAX:007788 |  |
| antibody | mouse PSD-95 primary antibody | Millipore MAB1598 | RRID:AB_11212185 | 1:500 |
| antibody | rabbit P-Ser73_PSD-95 primary antibody | Davids Biotechnologie | A061 | 1:12 |
| antibody | Donkey anti-Mouse IgG alexa fluor 555 | Invitrogen A31570 | RRID:AB_2536180 | 1:500 |
| antibody | Rabbit anti-mCherry | Abcam, ab167453 | RRID:AB_2571870 | 1:500 |
| antibody | Donkey anti-rabbit Alexa Fluor 555 | Invitrogen, A-31572 | RRID:AB_162543 | 1:500 |
| antibody | Donkey anti-rabbit Alexa Fluor 647 | Invitrogen, A31573 | RRID:AB_2536183 | 1:500 |
| recombinant DNA reagent | AAV1/2, *Camk2a*_PSD-95(WT):mCherry (PSD-95(WT)) | PMID: **32029829** |  | *Dlg4* cloned into Addgene plasmid # 114469 |
| recombinant DNA reagent | AAV1/2, *Camk2a*_PSD-95(S73A):mCherry (PSD-95(S73A)) | PMID: **32029829** |  | *Dlg4_S73A* cloned into Addgene plasmid # 114469 |
| recombinant DNA reagent | AAV1/2, *Camk2a*_mCherry (mCherry) | Addgene plasmid # 114469 | RRID:Addgene_114469 |  |
| software, algorithm | ImageJ (Fiji) | PMID: 22743772 | RRID:SCR_002285 |  |
| software, algorithm | Med-PC V Software Suite | Med Associate Inc. | SOF-736 | fear conditioning software |
| software, algorithm | Reconstruct | PMID: **15817063** | RRID:SCR_002716 |  |

## SUPPLEMENTARY FIGURES

**Supplementary Figure 1.**
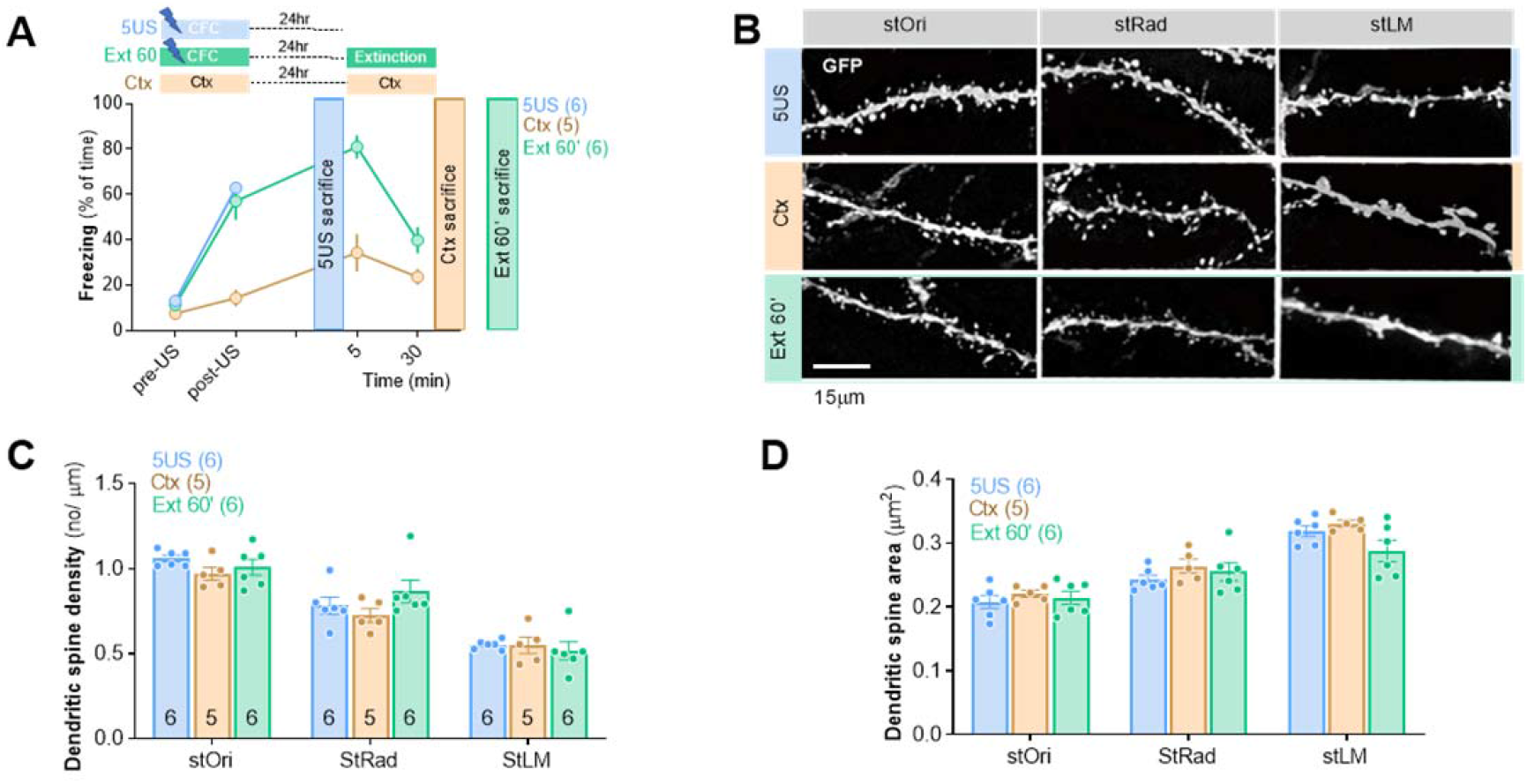
Synaptic plasticity induced by exposure to neutral context. Dendritic spines were analysed in three domains of dendritic tree of dCA1 area in Thy1-GFP(M) male mice: stOri, stRad and stLM. **(A)** Experimental timeline and freezing levels of mice from three experimental groups: 5US (mice sacrificed 1 day after CFC; n = 6), Ctx (mice sacrificed immediately after the second exposure to novel context, no foot shocks were delivered, n = 5) and Ext 60’ (mice sacrificed 60 minutes after contextual fear extinction session, n = 6). **(B)** Representative confocal images of dendrites (GFP) (maximum projections of z-stacks composed of 20 scans) are shown for three domains of the dendritic tree. **(C)** Summary of data showing dendritic spine density (repeated-measures ANOVA, effect of training: F(2, 14) = 1.620, P = 0.233). **(D)** Summary of data showing average dendritic spine area (repeated-measures ANOVA, effect of training: F(2, 14) = 3.162, P = 0.074). For C, D, each dot represents one mouse. For C means ± SEM are shown. For D, medians ± IQR are shown.

**Supplementary Figure 2.**
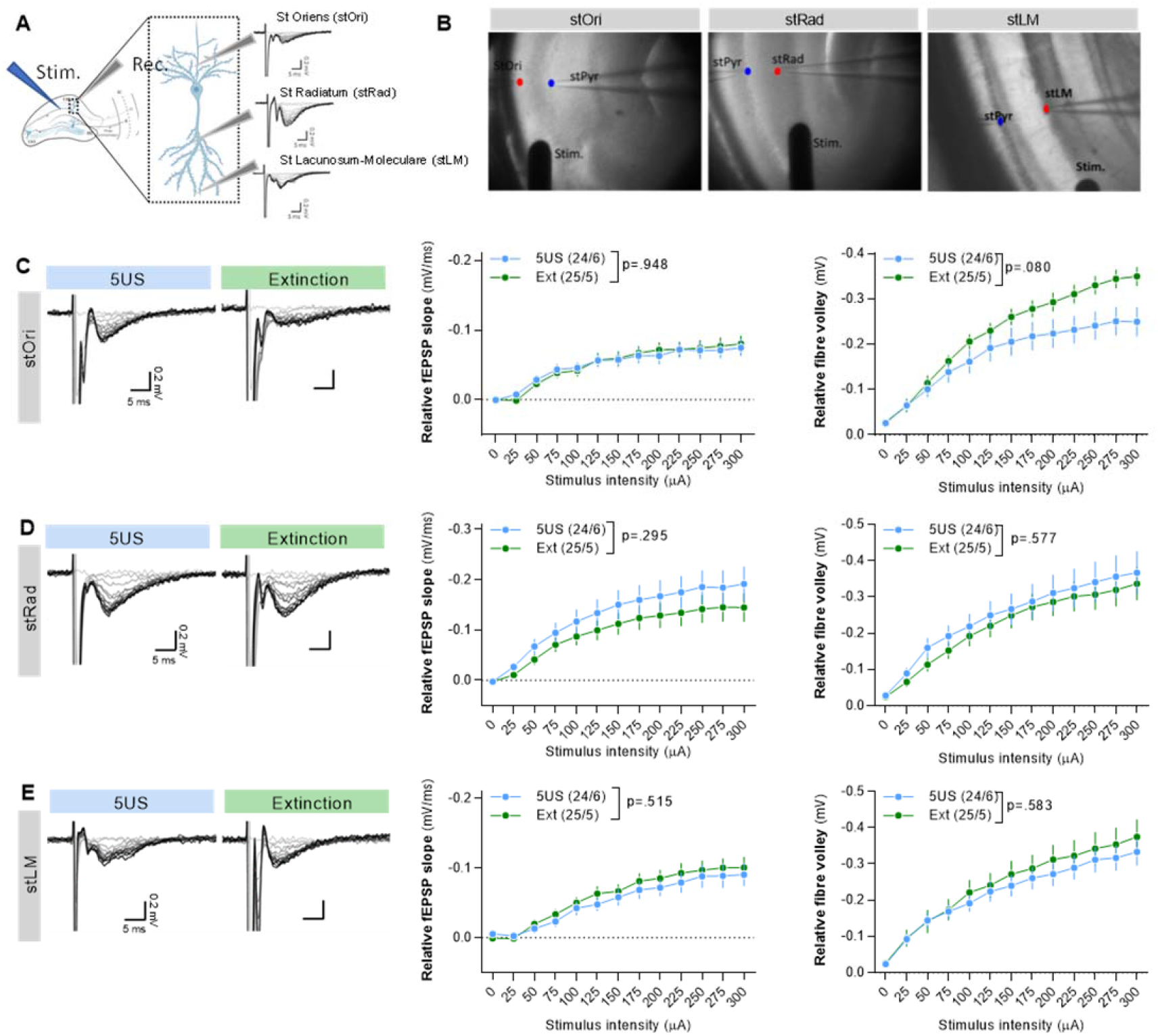
Synaptic plasticity induced in dCA1 during contextual fear extinction training is compensatory. **(A)** Experimental design. **(B)** Microphotographs of recording setups. Stim-stimulation electrodes (Stim.) and two recording electrodes were placed in stPyr (blue dot) and stOri, stRad or stLM (red dots). **(C-E) (left)** Representative fEPSPs evoked by stimuli of different intensities, **(middle)** input–output functions for stimulus intensity (repeated-measures ANOVA, effect of virus: stOri, F(1, 37) = 0.001, P = 0.971; stRad: F(1, 56) = 1.120, P = 0.294; stLM: F(1, 47) = 0.429, P = 0.515) and **(left)** fibre volley recorded in response to increasing intensities of stimulation (repeated-measures ANOVA, effect of virus: stOri, F(1, 43) = 3.198, P = 0.080; stRad: F(1, 47) = 0.314, P = 0.577; stLM: F(1, 44) = 0.305, P = 0.583). The numbers of the analysed sections/mice per experimental group are indicated in the legends. Means ± SEM are shown on the graphs.

**Supplementary Figure 3.**
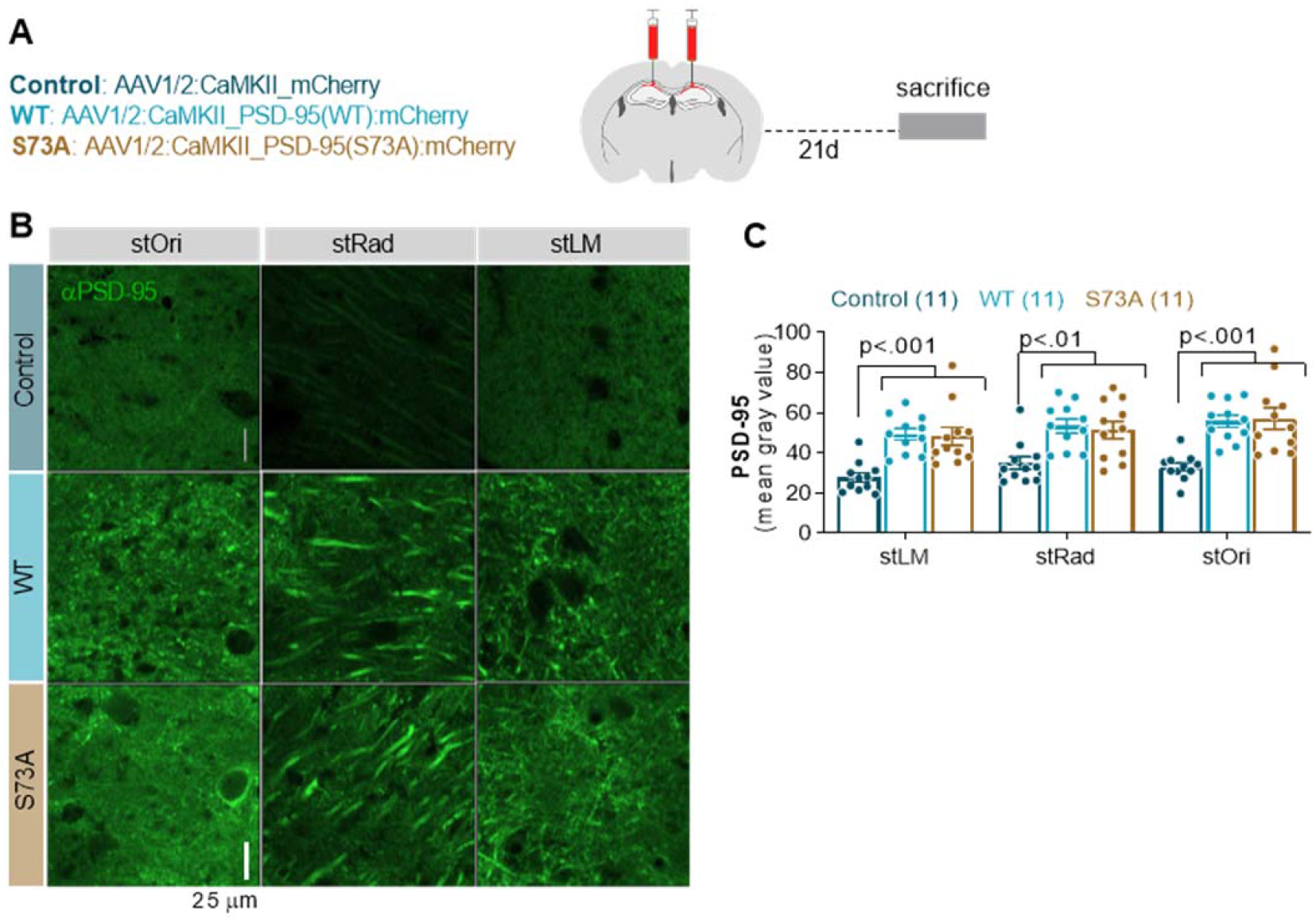
Validation of the viral vectors encoding PSD-95(WT) and PSD-95(S73A). **(A)** Experimental timeline. C57BL/6J male mice were stereotactically injected in the dCA1 with AAV1/2 encoding mCherry (Control, n=11) PSD-95(WT) (WT, n = 11) or PSD-95(S73A) (S73A, n = 11). Twenty one days later they were sacrificed. **(B)** Representative confocal scans of the PSD-95 immunostaining in dCA1 strata and **(C)** summary of data showing PSD-95 levels (two-way ANOVA with Tukey’s *post hoc* test, effect of virus: F(2, 30) = 13.1, P <0.001).

